# SARS-CoV-2 ORF6 disrupts innate immune signalling by inhibiting cellular mRNA export

**DOI:** 10.1101/2022.02.08.479664

**Authors:** Ross Hall, Anabel Guedán, Melvyn W. Yap, George R. Young, Ruth Harvey, Jonathan P. Stoye, Kate N. Bishop

## Abstract

SARS-CoV-2 is a betacoronavirus and the etiological agent of COVID-19, a devastating infectious disease. Due to its far-reaching effect on human health, there is an urgent and growing need to understand the viral molecular biology of SARS-CoV-2 and its interaction with the host cell. SARS-CoV-2 encodes 9 predicted accessory proteins, which are presumed to be dispensable for *in vitro* replication, most likely having a role in modulating the host cell environment to aid viral replication. Here we show that the ORF6 accessory protein interacts with cellular Rae1 to inhibit cellular protein production by blocking mRNA export. We utilised cell fractionation coupled with mRNAseq to explore which cellular mRNA species are affected by ORF6 expression and show that ORF6 can inhibit the export of many mRNA including those encoding antiviral factors such as IRF1 and RIG-I. We also show that export of these mRNA is blocked in the context of SARS-CoV-2 infection. Together, our studies identify a novel mechanism by which SARS-CoV-2 can manipulate the host cell environment to supress antiviral responses, providing further understanding to the replication strategies of a virus that has caused an unprecedented global health crisis.

**Author Summary:** SARS-CoV-2 is the virus responsible for the current COVID-19 pandemic. Coronaviruses, like SARS-CoV-2, replicate their genome in the cytoplasm of the host cell by hijacking the cellular machinery. In addition to structural proteins and viral enzymes, SARS-CoV-2 encodes 9 accessory proteins. Although these are not required for in vitro replication, they are thought to modulate the host cell environment to favour viral replication. In this work, we show that the ORF6 accessory protein can supress cellular protein production by blocking mRNA nuclear export through interacting with the cellular protein Rae1, a known mRNA export factor. We also investigated which cellular mRNAs were retained in the nucleus when ORF6 was overexpressed. Interestingly, we found that ORF6 inhibited the export of many different mRNAs, including those encoding antiviral factors, like IRF1 and RIG-I, even in the absence of stimulation by interferon. Importantly, we found that the export of these mRNAs was similarly affected in the context of SARS-CoV-2 infection. Therefore, we believe we have identified a novel mechanism that SARS-CoV-2 uses to suppress antiviral responses in order to make the cell more permissive to infection.

## Introduction

Severe acute respiratory syndrome coronavirus 2 (SARS-CoV-2) is an enveloped single-stranded, positive-sense RNA virus of the *Betacoronovirus* genus and is the etiological agent of the highly infectious disease, COVID-19. SARS-CoV-2 replicates in the epithelial cells of the respiratory tract causing extensive respiratory symptoms, with severe cases leading to mortality [1-4]. SARS-CoV-2 expresses four structural proteins, spike (S), membrane (M), envelope (E) and nucleoprotein (N) and 16 non-structural proteins (nsps), as well as nine predicted accessory proteins; ORF3a, ORF3b, ORF6, ORF7a, ORF7b, ORF8, ORF9b, ORF9c and ORF10 [2, 5]. These accessory proteins are presumed dispensable for *in vitro* replication and have been suggested to play a role in pathogenicity and host cell modulation [6, 7]. Little is known about the function of these accessory proteins and how they affect the cellular environment to facilitate efficient SARS-CoV-2 replication.

SARS-CoV-2 shares a high degree of homology with the etiological agent of the 2003 SARS outbreak, SARS-CoV (hereafter known as SARS-CoV-1). Indeed, the accessory proteins of these two betacoronaviruses have high homology, with ORF3b and ORF6 being the most divergent. ORF6 from SARS-CoV-1 is a 7.3 kDa protein, which has been reported to inhibit IFN signalling by disrupting the nuclear import of STAT1/STAT2 and IRF3 [8, 9]. This is dependent on ORF6 interacting with the cellular Rae1-Nup98 complex [10], which spans the nuclear envelope, providing a gateway between the nucleus and cytoplasm, and helps shuttle proteins and mRNA between the two compartments [11, 12]. Other viruses have also been shown to disrupt this nuclear export pathway to aid viral propagation. VSV matrix (M) and herpesvirus ORF10 both inhibit mRNA nuclear export by disrupting Nup98-Rae1 function, while Influenza virus NS1 renders cells more permissive to replication by downregulating Nup98 expression thereby inhibiting Rae1-dependent mRNA nuclear export [13-16]. As coronaviruses transcribe their genome in the cytoplasm, they are not dependent on nuclear export function for viral replication. Indeed, NSP1 proteins from coronaviruses have been shown to inhibit the NXF1 mRNA export pathway, potentially disrupting the expression of a range of cellular proteins [17].

Given the potential importance of the SARS-CoV-2 accessory proteins we set out to investigate their functions. Strikingly, we found that ORF6 inhibited protein production and we show that in addition to disrupting STAT1/2 import, ORF6 can also block the export of cellular mRNA from the nucleus into the cytoplasm, effectively blocking protein translation. In agreement with other very recent studies of SARS-CoV-2 ORF6, this is also dependent on interaction with the Nup98-Rae1 complex [10, 18, 19]. Here, we utilise cellular fractionation and mRNAseq to identify the mRNA species blocked by ORF6. We show that ORF6 inhibits export of a broad range of mRNAs, in particular several interferon-upregulated genes that encode antiviral factors, including RIG-I and IRF1. This shows that SARS-CoV-2 ORF6 can disrupt host cell innate immune signalling by blocking mRNA nuclear export and suggests that ORF6 may function to inhibit the earliest stages of innate signalling by downregulating expression of pathogen recognition receptors (PRRs) and antiviral transcription factors, such as IRF1.

## Results

### SARS-CoV-2 ORF6 Inhibits protein expression

To investigate the role of SARS-CoV-2 accessory proteins, we first generated a panel of HIV-1 virus-like-particles (VLPs) encoding each open-reading frame (ORF) to transduce a range of different cell lines to express each accessory protein individually. VLPs were synthesised by co-transfecting HEK293T cells with plasmids encoding HIV-1 Gag-Pol (pCMVΔR8.91), VSV-G (pVSV-G) and a pLVX-StrepII-ORF vector, encoding either ORF3a, ORF3b, ORF6, ORF7a, ORF7b, ORF8, ORF9b, ORF9c or ORF10. VLPs titres were determined after 72 hours and at least 10^4 infectious units/ml were detected for all VLPs except for VLP encoding ORF6 which had undetectable infectious units (Fig 1A, upper panel). To determine if this was due to decreased infectivity or decreased particle production, transfected HEK293T cell lysates were analysed by immunoblotting for the HIV-1 structural polyprotein, Gag. Gag was detectable in all cells, except for cells transfected with pLVX-StrepII-ORF6 (Fig 1A, lower panel). This suggested that ORF6 expression resulted in inhibition of Gag production preventing formation of VLPs.

**Fig 1.**
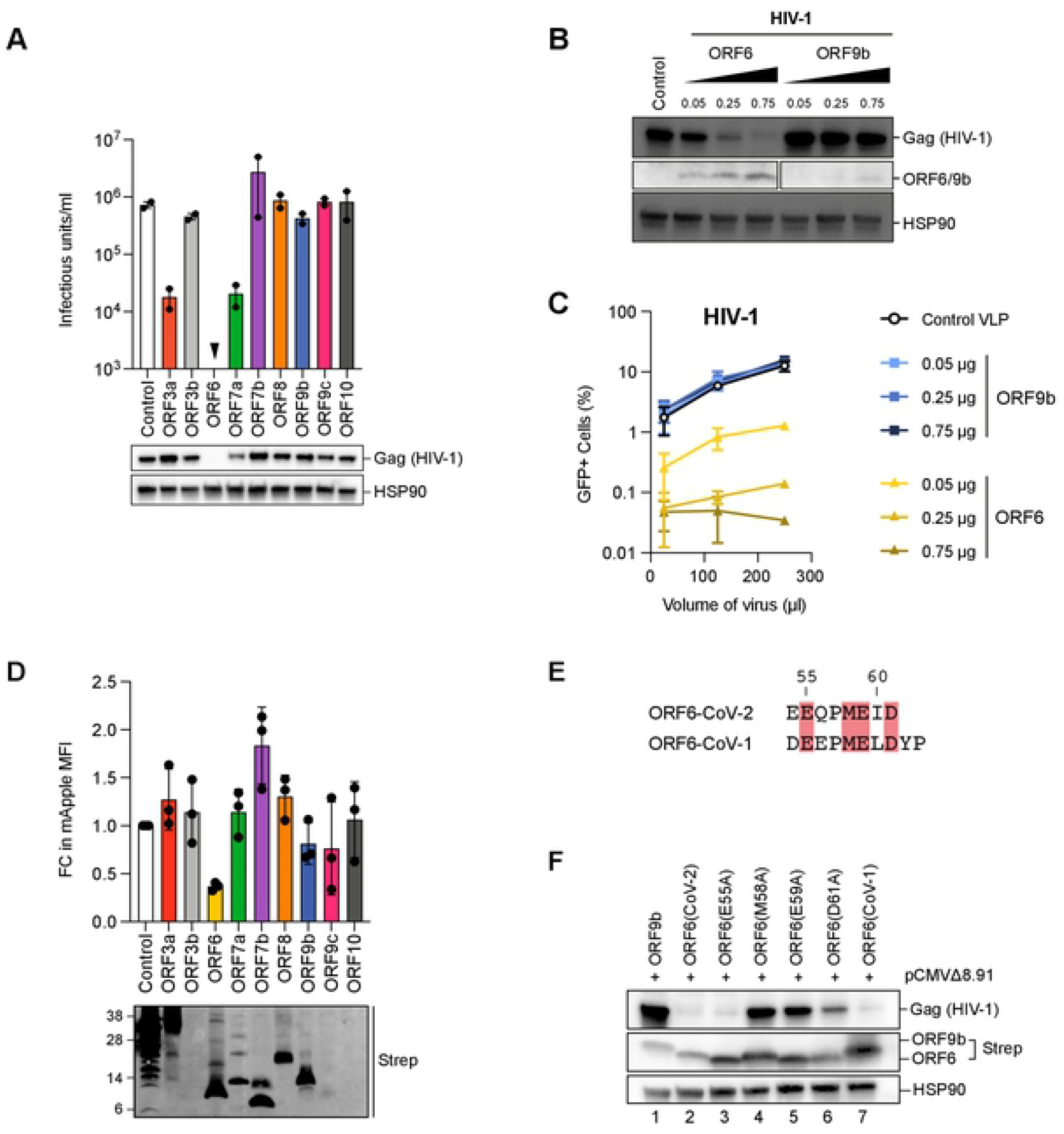
SARS-CoV-2 ORF6 inhibits protein expression. (A) To generate VLPs, HEK293T cells were co-transfected with plasmids encoding HIV-1 Gag-Pol, VSV-G and a pLVX-StrepII-ORF vector encoding either SARS-CoV-2 ORF3a, ORF3b, ORF6, ORF7a, ORF7b, ORF8, ORF9b, ORF9c or ORF10 or an empty vector control. VLPs were harvested and titrated in HeLa cells. The infectious units/ml are shown in the bar graph. Points indicated independent biological repeats. Transfected cell lysates were separated by SDS-PAGE and analysed for Gag(HIV-1) and HSP90 by immunoblotting. (B,C) HEK293T cells were transfected with plasmids encoding HIV-1 Gag-Pol, VSV-G and either increasing amounts of pLVX-StrepII-ORF6 or pLVX-StrepII-ORF9b as well as a GFP reporter plasmid, pCSGW. (B) Transfected cell lysates were separated by SDS-PAGE and analysed for Gag(HIV-1), ORF6 or ORF9b and HSP90 by immunoblotting. (C) HeLa cells were infected with increasing amounts of HIV-1 VLPs. Three days post infection, the percentage of GFP positive cells was determined by flow cytometry. Graph shows the mean and range of two biological repeats. (D) HEK293T cells were transfected with the pLVX-EF1α plasmid encoding strep-tagged GFP(Control), ORF3a, ORF3b, ORF6, ORF7a, ORF7b, ORF8, ORF9b, ORF9c or ORF10 and a plasmid encoding mApple. After 48h, cells were fixed and the mApple median fluorescent intensity (MFI) measured by flow cytometry. The fold change (FC) in MFI is plotted relative to the control for independent biological repeats. Error bars show SEM. Transfected cell lysates were analysed by SDS-PAGE and immunoblotting with anti-strep. (E) Amino acid alignment of the C-terminus region of ORF6 from SARS-CoV-2 and SARS-CoV-1 with conserved residues highlighted in red. (F) HEK293T cells were transfected with pLVX-EF1α encoding strep-tagged ORF9b, ORF6(CoV-2), ORF6(CoV-1) or indicated ORF6(CoV-2) mutant and a plasmid encoding HIV-1 Gag-Pol. After 24h, cell lysates were separated by SDS-PAGE and analysed for Gag(HIV-1), Strep and HSP90 by immunoblotting.

To determine if there was a dose-dependent effect on Gag expression and VLP production, we next generated GFP-reporter HIV-1 VLPs in the presence of increasing amounts of ORF6 or ORF9b as a control. Transfected HEK293T cell lysates were analysed by immunoblotting for HIV-1 Gag and ORF6/ORF9b. Gag expression was reduced in a dose-dependent manner only with increasing amounts of ORF6 (Fig 1B). VLPs generated from these transfections were used to transduce HeLa cells and the percentage of cells expressing GFP was determined after 72 hours by flow cytometry to measure VLP titre. As with Gag expression, HIV-1 VLP titres were severely reduced in a dose dependent manner compared to the control when produced in the presence of ORF6 (Fig 1C, yellow lines) but not ORF9b (Fig 1C, blue lines).

To determine if ORF6 inhibition was specific to HIV-1 Gag expression, we next examined the effect of ORF6 on the expression of MLV Gag and mApple fluorescent protein, which have completely different protein sequences to HIV-1 Gag. As with HIV-1 Gag, ORF6 inhibited MLV Gag expression and VLP production in a dose-dependent manner (S1A and B Figs). Furthermore, when HEK293T cells were co-transfected with plasmids expressing mApple and each SARS-CoV-2 ORF and the median fluorescent intensity (MFI) of mApple was measured by flow cytometry, ORF6 reduced the MFI approximately 3-fold compared to the control (Fig 1D, upper panel). None of the other ORFs had an effect on the MFI of mApple (Fig 1D, upper panel), although immunoblotting revealed very low expression of ORF3b and ORF10 (Fig 1D, lower panel), which has also been observed by others [5]. Taken together our results suggest that ORF6 can inhibit expression of multiple proteins expressed from a vector.

Previous work has shown that the C-terminal region of ORF6 is important for inhibiting IRF3 and STAT nuclear import [5, 8, 10]. To determine if this region is also important for ORF6-dependent inhibition of protein expression, we introduced alanine substitutions at residues in the C-terminus that were conserved between SARS-CoV-2 and SARS-CoV-1 ORF6 proteins, E55A, M58A, E59A or D61A (Fig 1E, highlighted in red). HEK293T cells were co-transfected with plasmids encoding each mutant ORF6 or wild type ORF6 from either SARS-CoV-2 or SARS-CoV-1 together with HIV-1 Gag and cell lysates were analysed by immunoblotting (Fig 1F). As before, ORF6(CoV-2) (Fig 1F, lane 2) reduced Gag expression compared to the ORF9b control (lane 1). In addition, ORF6(CoV-1) (lane 7) and mutant E55A (lane 3) were also able to inhibit Gag expression. However, ORF6 mutants M58A (lane 4), E59A (lane 5) and D61A (lane 6) had a much weaker effect on Gag expression suggesting that residues M58, E59 and D61 are essential for reducing protein expression. ORF6 mutants M58A, E59A and D61A also had a much weaker effect on mApple expression compared to wild type ORF6 (Fig S1C).

### ORF6 inhibits protein expression by interfering with Rae1

There are several ways that ORF6 could block protein expression, for example inhibiting transcription, mRNA nuclear export or translation. Biochemical studies have indicated that SARS-CoV-2 ORF6 interacts with the Nup98-Rae1 complex [5] which is important for mRNA nuclear export. Specifically, Rae1 helps shuttle nascent mRNA into the cytoplasm for translation [11, 12, 20]. Therefore, to investigate whether ORF6 could block protein expression by interfering with the Nup98-Rae1 complex, we overexpressed Rae1 in HEK293T cells together with HIV-1 Gag and ORF6. Fig 2A shows that expressing exogenous Rae1 partially rescued Gag expression in the presence of ORF6 (Fig 2A, compare lanes 2 and 3). Relief from ORF6 inhibition could be further enhanced by introduction of a mutation known to affect the inhibitory interaction between the VSV M protein and Rae1 [13]. Whilst expression of Rae1(T306I) had a similar effect to overexpressing wild type Rae1 (lane 5), expression of Rae1(R305G) fully rescued Gag expression to the levels seen in the ORF9b control (compare lane 4 with lane 1). This suggests that ORF6 interacts with Rae1 to inhibit protein expression and that R305 is important for the interaction.

**Fig 2.**
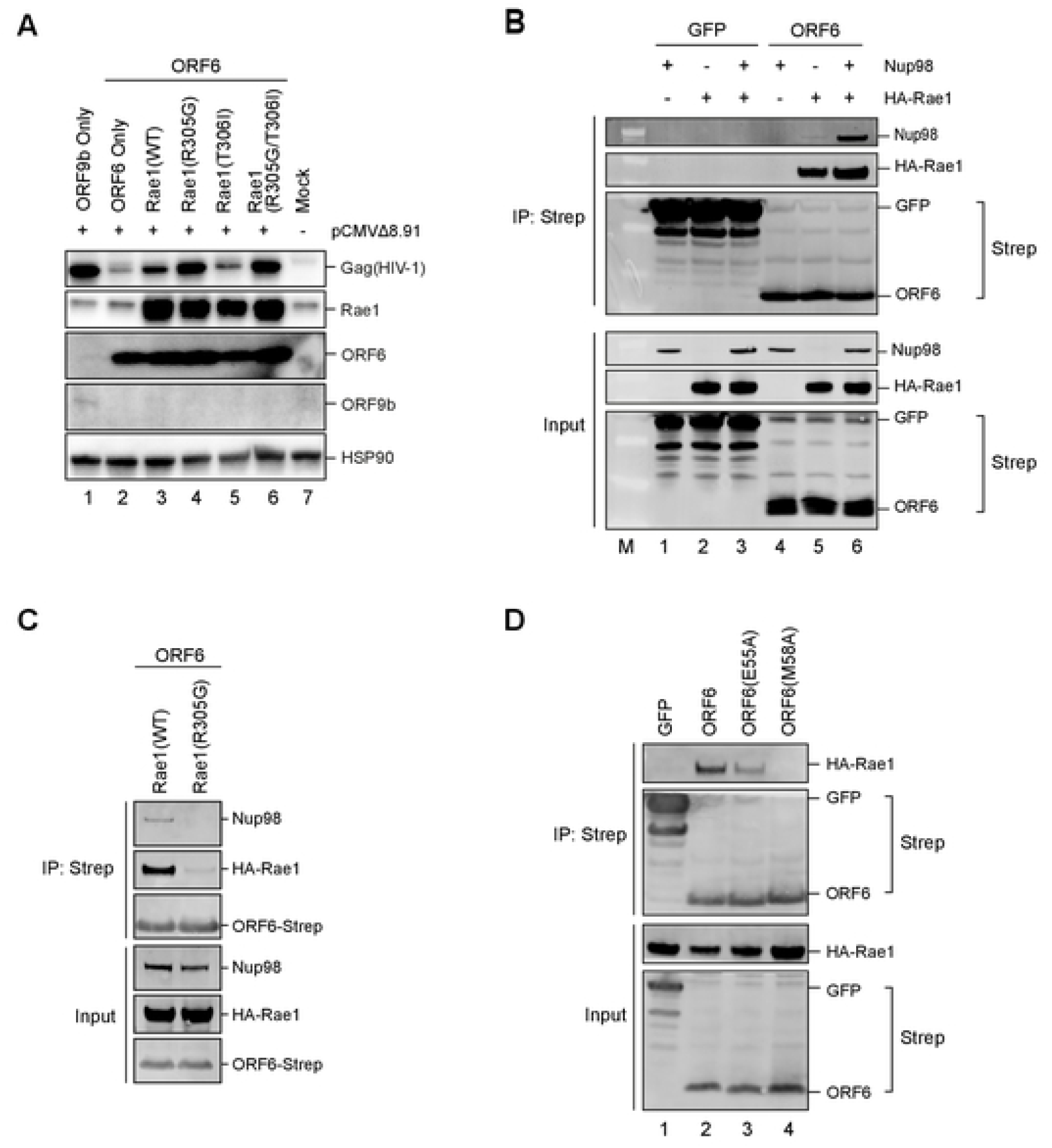
ORF6 inhibits protein expression by interfering with Rae1. (A) HEK293T cells were co-transfected with plasmids encoding HIV-1 Gag-Pol, ORF6 or ORF9b and Rae1 or indicated Rae1 mutant. Mock cells were untransfected. After 48h, cell lysates were separated by SDS-PAGE and analysed for Gag(HIV-1), Rae1, ORF6 or ORF9b and HSP90 by immunoblotting. (B) HEK293T cells were transfected with plasmids encoding Twin-Strep-tagged GFP or ORF6 and Nup98 and/or HA-Rae1. After 24h, Twin-Strep-tagged proteins were immunoprecipitated with MagStrep beads and proteins eluted with biotin. Input lysates and eluate were separated by SDS-PAGE and analysed by immunoblotting for Strep, Nup98 and HA. (C) HEK293T cells were transfected with plasmids encoding Twin-Strep-tagged GFP or ORF6, Nup98 and either HA-Rae1(WT) or HA-Rae1(R305G). After 24h, Twin-Strep-tagged proteins were immunoprecipitated and analysed as in (C). (D) HEK293T cells were transfected with plasmids encoding Twin-Strep-tagged proteins; ORF6(CoV-2), ORF6(E55A), ORF6(M58A) or GFP and HA-Rae1(WT). After 24h, Twin-Strep-tagged proteins were immunoprecipitated with MagStep beads and input lysates and eluates separated by SDS-PAGE and analysed by immunoblotting for Strep and HA.

To confirm the interaction of ORF6 with Rae1 and examine the interaction with Nup98, we performed co-immunoprecipitations (co-IP). HEK293T cells were co-transfected with plasmids expressing Strep-tagged GFP or ORF6 with Nup98 and/or HA-Rae1. After 48 hrs, cells were lysed and GFP or ORF6 immunoprecipitated with anti-Strep antibody. Input lysates and precipitated proteins were analysed by immunoblotting for Strep, HA and Nup98. Both GFP and ORF6 were detected in the eluate, confirming successful immunoprecipitation (Fig 2B). The GFP control did not co-immunoprecipitate with either HA-Rae1 or Nup98. However, HA-Rae1 was detectable in the eluate when co-expressed with ORF6, confirming an interaction between these two proteins. Rae1 was detectable whether Nup98 was co-expressed or not, suggesting that the ORF6 interaction with Rae1 is not dependent on Nup98. Conversely, Nup98 was only detectable in the eluate when co-expressed with both ORF6 and Rae1. ORF6 and Rae1 also co-immunoprecipitated with a small amount of endogenous Nup98 (Fig 2B, lane 5). This suggests that the interaction between Nup98 and ORF6 is not direct but is instead dependent on Rae1. Importantly, the Rae1(R305G) mutation that conferred resistance to ORF6-dependent inhibition of protein expression (Fig 2A) did not co-immunoprecipitate with ORF6 (Fig 2C) and Nup98 was also not present under these conditions, confirming that Nup98 interacts with ORF6 via Rae1. We also tested whether the ORF6 mutants could co-precipitate Rae1 (Fig 2D). As expected, ORF6 and ORF6(E55A), which both inhibited Gag expression (Fig 1F), co-immunoprecipitated with Rae1 whilst ORF6(M58A) did not (Fig 2D). Additionally, we also observed that ORF6(CoV-1) co-immunoprecipitated with Rae1 (S2A Fig), and that both ORF6 homologues co-immunoprecipitated with endogenous Rae1 (S2B Fig). Overall, this implies that the effect of ORF6 on protein expression is mediated through an interaction with Rae1, suggesting that the block is to mRNA export.

### ORF6 inhibits the nuclear export of cellular mRNA

To investigate the effect of ORF6 on mRNA export directly, we first set up an assay to separate nuclear (Nucl) and cytoplasmic (Cyto) fractions and measure mRNA levels in each by qPCR. A higher ratio of Nucl:Cyto mRNA compared to the control would suggest that the mRNA is being retained in the nucleus. As a proof-of-principle, we treated cells with Leptomycin B (LMB), which inhibits Chromosomal Maintenance 1 (CRM1)-dependent mRNA export [21] (S3 Fig). HEK293T cells were transfected with a plasmid expressing GFP, treated with LMB for 16 hrs and then fractionated. LMB clearly reduced GFP expression in cells (S3A and B Fig). Absolute levels of GFP mRNA were determined by qPCR in the Nucl and Cyto fractions (S3C Fig). Although total GFP RNA levels were reduced, LMB-treated cells also had a higher Nucl:Cyto GFP mRNA ratio compared to untreated cells, at 0.93 compared to 0.35 (S3D Fig). Like LMB-treated cells, cells expressing ORF6 also reduced GFP expression (S3E and F Fig) and increased the ratio of Nucl:Cyto GFP mRNA (S3G and H Fig), while cells expressing ORF6(M58A) had a similar Nucl:Cyto GFP mRNA as the control cells expressing ORF9b. Overall, this indicates that ORF6 inhibits nuclear export of GFP mRNA, in a Rae1-dependent manner.

So far, we have shown that ORF6 can inhibit exogenous protein mRNA nuclear export and that this likely accounts for the block to protein production. To investigate whether ORF6 also inhibits endogenous cellular mRNA export, and to determine whether specific proteins are affected, we performed mRNAseq on nuclear and cytoplasmic fractions of HEK293T cells either untransfected (mock) or transfected with ORF6 or GFP. Transfections were performed in duplicate and after 24 hrs, one sample for each transfection was treated with IFN-β (1,000 units/ml) for 16 hrs. Cells were then fractionated and a sample was taken for protein analysis. RNA was extracted from the remaining fraction and mRNA libraries were prepared and sequenced (Fig 3A). Lysates were analysed by immunoblotting with anti-Strep antibodies to confirm ORF6/GFP expression, HSP90 (cytoplasmic marker) and lamin B1 (nuclear marker), to confirm that cellular fractionation was successful and phosphorylated STAT1 (pSTAT-Y701) to confirm activation of the type-I IFN signalling cascade (S4A Fig).

**Fig 3.**
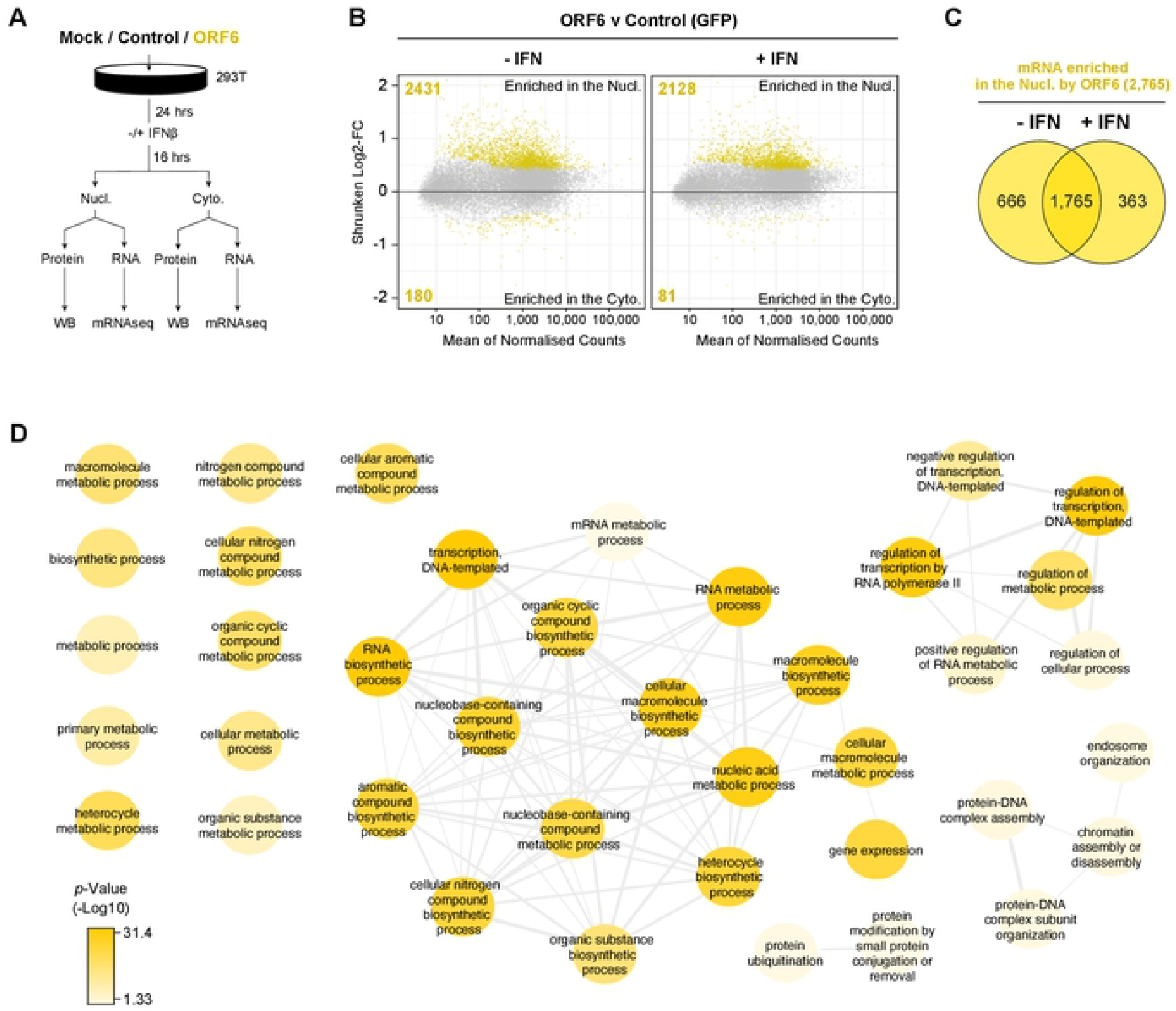
ORF6 inhibits the nuclear export of cellular mRNA. HEK293T cells were transfected with pLVX-EF1α-GFP or pLVX-EF1α-ORF6 or left untransfected (mock). After 24h, cells were either treated with IFN-β (1,000 units/ml) for 16 hrs or left untreated. Cells were then harvested and fractionated into nuclear (Nucl) and cytoplasmic (Cyto) fractions. Cells were either lysed for immunoblot analysis (see S4A Fig) or RNA was extracted for mRNAseq. (A) Schematic of experiment design. (B) The log2-fold change (Log2-FC) in mRNA abundance was calculated between the Nucl and Cyto fractions for both GFP and ORF6 expressing cells, without and with IFN treatment (see S4B Fig). The log2-FC in mRNA abundance was then directly compared between ORF6 and GFP, without (left panel) and with (right panel) IFN, to determine which mRNAs were specifically enriched in the nucleus or cytoplasm by ORF6. The log-2FC was weighted against the adjusted *p*-value (shrunken Log2-FC) to show significantly enriched mRNAs (yellow points). The number of mRNAs significantly enriched is shown. The mRNA count was normalised and averaged between the three biological repeats. (C) Venn diagram of the number of mRNAs that were significantly enriched in the nucleus by ORF6 with and without IFN. (D) mRNAs significantly enriched by ORF6 without IFN treatment were analysed for enriched GO terms, which were filtered and mapped using the REVIGO tool. Highly similar GO terms are grouped and linked. Colour intensity reflects significance levels.

Next, we determined the distribution of mRNA between the cytoplasm and nucleus during ORF6 and GFP expression without and with IFN-β treatment compared to the untransfected cells (S4B Fig). ORF6 was then further compared to GFP to determine which mRNA species were specifically enriched in the nucleus or cytoplasm in the presence of ORF6 (Fig 3B). A positive fold change (FC) indicates that an mRNA species is enriched in the nucleus, while a negative FC suggests that an mRNA species is enriched in the cytoplasm. In total, in the absence of IFN, 2,431 mRNA species were enriched in the nucleus with ORF6, while only 180 mRNA species were enriched in the cytoplasm, compared to the control (Fig 3B, left panel, highlighted in yellow). With IFN-β treatment, 2,128 mRNA species were enriched in the nucleus with ORF6, while 81 were enriched in the cytoplasm, compared to the control (Fig 3B, right panel). There was considerable overlap (1,765 mRNA) between the mRNA species enriched in the nucleus in both conditions (Fig 3C). These genes were found to include a wide range of Gene Ontology (GO) terms (Fig 3D), highlighting their breadth of function. These data suggest that ORF6 not only inhibits mRNA export of exogenous mRNA such as GFP but, more importantly, also inhibits the nuclear export of a broad range of cellular mRNA.

### ORF6 inhibits the nuclear export of IFN-upregulated mRNA

As many mRNA were retained in the nucleus, we decided to investigate whether genes with potential anti-viral functions were among them. Firstly, we identified potential ISGs in our dataset by determining which mRNA species were upregulated by IFN-β treatment in cells transfected with our control plasmid (S5A Fig). We found 134 mRNAs species that were significantly upregulated in either the cytoplasm and/or nucleus after IFN-β treatment and labelled them interferon upregulated genes (IUGs) (Fig 4A and S5B Fig). Importantly, these IUGs encoded many proteins with roles in inhibiting viral replication and immune regulation (S5C Fig) and included well characterised ISGs like BST2 and IFITM1. Next, as ORF6 has been reported to inhibit IFN signalling, we measured mRNA expression in the presence of ORF6 compared to control cells in the context of IFN-β treatment to determine if the expression of any of these identified IUGs was modulated by ORF6. Interestingly, of the 134 IUGs we identified, 69 IUGs were significantly downregulated in the cytoplasm in the presence of ORF6 (yellow tones) and 2 were significantly upregulated (dark blue tones) (Fig 4A, top panel). In the nuclear fraction, 62 IUGs were significantly downregulated in the presence of ORF6 while 10 were significantly upregulated (Fig 4A, bottom panel). In total, 79 unique IUGs were significantly downregulated by ORF6, while 11 unique IUGs were upregulated (S5D Fig). This supports the notion that ORF6 can inhibit IUG mRNA expression. However, in addition to the IUGs, ORF6 modulated the expression of many other mRNA compared to the control. S5E Fig summarises the total number of mRNA species upregulated or downregulated in the nucleus and/or cytoplasm by ORF6 in IFN treated and untreated cells.

**Fig 4.**
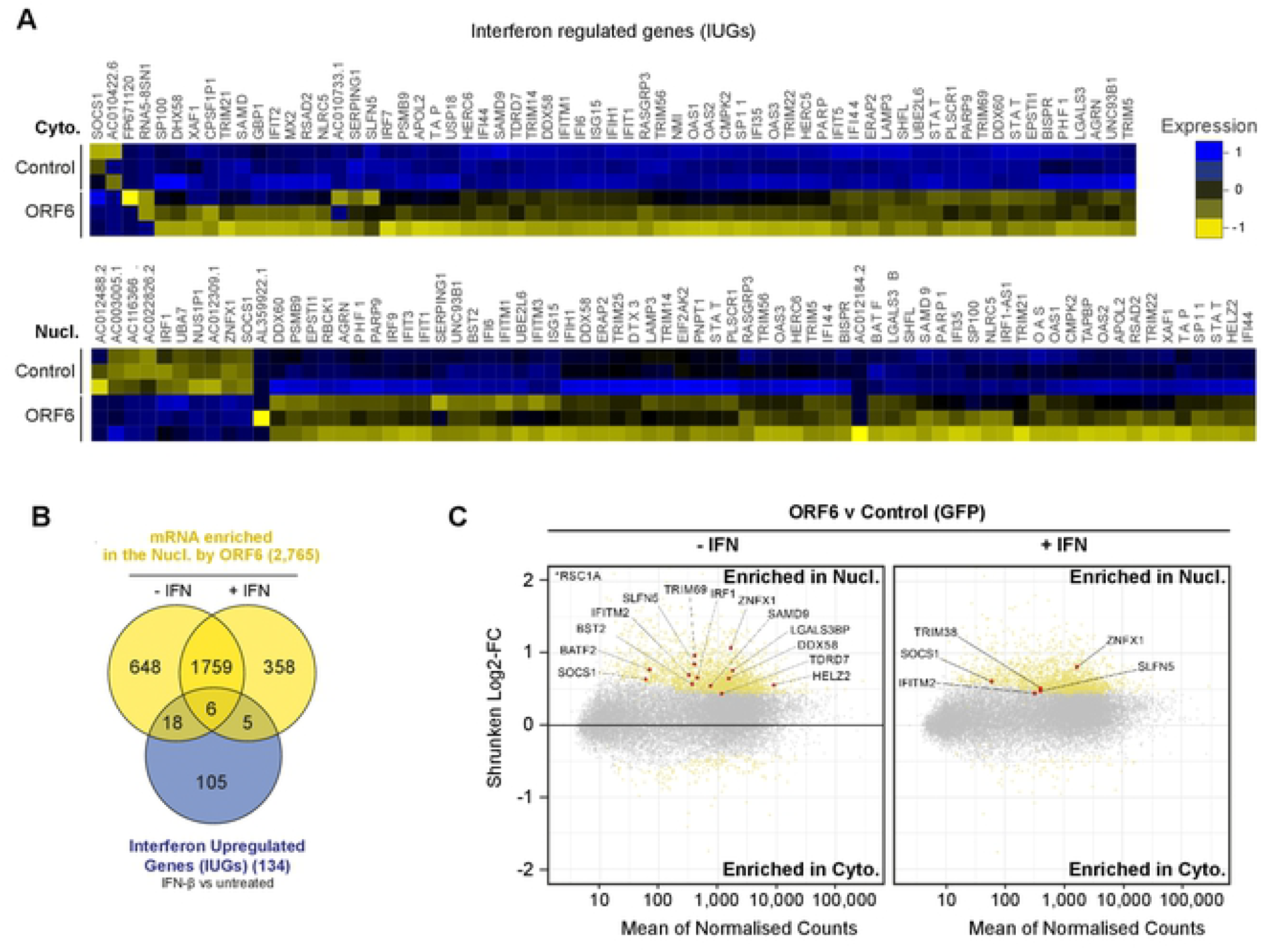
ORF6 inhibits the nuclear export of IFN-upregulated mRNA. Interferon upregulated genes (IUGs) were identified as mRNAs that were significantly upregulated after IFN treatment compared to untreated cells (see S5A Fig). These IUGs were then compared to the list of mRNAs that were significantly upregulated or downregulated by ORF6 compared to the GFP control (see S5E Fig), in either the Cyto (upper panel) or Nucl (lower panel) fractions. The heat maps show the relative Log2-Fold Change in IUG expression in cells expressing GFP (Control) or ORF6 after IFN treatment, for three biological repeats. Venn diagram showing the number of IUG mRNAs that were significantly enriched in the nucleus by ORF6 with and without IFN treatment. (C) The log2-fold change (Log2-FC) in mRNA abundance was calculated between the Nucl and Cyto fractions for both GFP and ORF6 expressing cells, without and with IFN treatment, as in Fig 3B, to determine which mRNA were specifically enriched by ORF6. The log-2FC was weighted against the adjusted *p*-value (shrunken Log2-FC) to show significantly enriched mRNA (yellow points). IUGs are highlighted in red and labelled.

Next, we further investigated the effect of ORF6 on the IUGs we had identified. Intriguingly, of the 2,431 mRNA species that were enriched in the nucleus by ORF6 in the absence of IFN treatment (Fig 3B), 24 were identified as IUGs (Fig 4B), including the antiviral factors TRIM69, BST2, ZNFX1, IRF1, and DDX58 (RIG-I) (Fig 4C, left panel). This suggests that ORF6 blocks the mRNA export of IUGs even when they are expressed at a basal level, which has significant implications for the antiviral state of the host cell. Furthermore, after IFN stimulation, ORF6 enriched 11 IUGs in the nucleus including TRIM38, SOCS1, IFITM2, SLFN5 and ZNFX1 (Fig 4C, right panel). Six of the IUGs were enriched in the nucleus in both IFN-treated and untreated cells (Fig 4B). Altogether, we identified 29 IUGs that were retained in the nucleus in the presence of ORF6 (Fig 4B) suggesting that in addition to inhibiting ISG expression, ORF6 can disrupt innate signalling by preventing the nuclear export of mRNA species that encode key antiviral factors that are both basally expressed and induced by IFN-β, like RIG-I, IRF1 and ZNFX1.

### SARS-CoV-2 inhibits the export of cellular mRNA

Our results demonstrate that expression of ORF6 in cells inhibits mRNA nuclear export with the effect of inhibiting protein expression. However, in the context of a viral infection, another viral protein may counteract the actions of ORF6. Therefore, to investigate whether mRNA export is also blocked during a SARS-CoV-2 infection, we analysed the nuclear accumulation of specific mRNAs in infected cells. Vero cells were infected with one of four SARS-CoV-2 variants: Eng/2, Alpha, Beta or Delta at a high MOI for 48 hrs, after which cells were either fixed, permeabilised and probed with an anti-ORF6 antibody or separated into Nucl and Cyto fractions and processed for protein and RNA extraction. For all infections, ORF6 predominantly localised to the cytoplasm with some perinuclear staining (Fig 5A). The pattern of ORF6 expression was less diffuse in SARS-CoV-2(delta) infected cells than the other infections. We also saw some co-localisation of ORF6 with Nup98 (S6A Fig). Immunoblot analysis of cell lysates for HSP90 (cytoplasmic) and Histone H3 (nuclear) confirmed that fractionation of samples was successful (S6B Fig). ORF6 was also detected predominately in the cytoplasm for most infections, although in two independent infections with SARS-CoV-2(delta) there was more ORF6 in the nuclear fraction (S6B and D Fig). This was probably linked to the poor viability of SARS-CoV-2(delta)-infected cells. In order to investigate mRNA export, we chose eight genes to analyse from our mRNAseq data: We identified three mRNAs that were most enriched in the nucleus in the presence of ORF6 (FAM222A, SPOCK1, SERTAD1, together classed “non-antiviral”) (S6E Fig, red dots) as well as four mRNA encoding antiviral IUGs that were significantly enriched in the nucleus (RIG-I, IRF1, ZNFX1 and TRIM38) (Fig 4B, red dots). GAPDH was not significantly enriched in the nucleus with ORF6 and so was used as a control. Overall, all four SARS-CoV-2 variants increased the average Nucl:Cyto ratio of FAM222A, SPOCK1 and SERTAD1 mRNAs compared to uninfected cells (Fig 5B) while the Nucl:Cyto ratio of GAPDH mRNA remained constant. This suggests that ORF6 can block the export of these mRNA species in the context of SARS-CoV-2 infection. Strikingly, there was a large increase in the average Nucl:Cyto ratio of both RIG-I and IRF1 mRNA after infection with all four variants (Fig 5B). ZNFX1 was also enriched in the nucleus for all infections except SARS-CoV-2 (delta). However, TRIM38, which was only enriched in the nucleus by ORF6 after IFN treatment (Fig 4B, right panel), was only marginally enriched in the nucleus after infection. Overall, we observed that SARS-CoV-2 infection induced retention of mRNA in the nucleus similar to ORF6 expression alone and that export of mRNAs involved in transcription (SERTAD1) and immune regulation (RIG-I and IRF1) was inhibited, suggesting that ORF6 has a broad effect on cellular pathways during infection.

**Fig 5.**
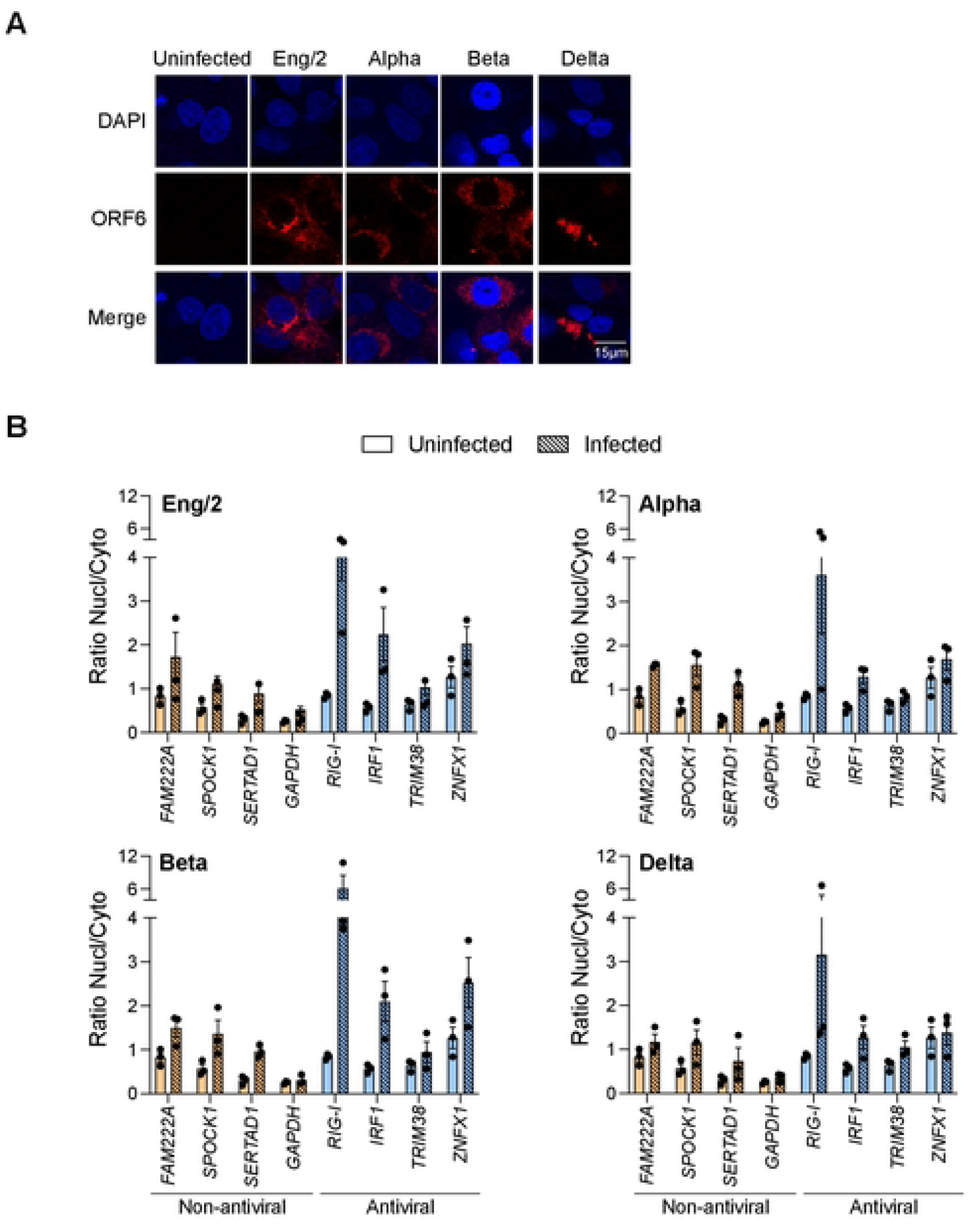
SARS-CoV-2 inhibits the export of cellular mRNA. Vero cells were inoculated with four SARS-CoV-2 variants (Eng/2, Alpha, Beta, Delta) at a MOI ≥ 1 for 48h. (A) Cells were fixed and analysed by immunofluorescence with anti-ORF6 and DAPI. (B) Cells were fractionated, RNA was extracted, and cDNA was generated. mRNA levels of four non-antiviral (orange) and four antiviral (blue) proteins in both Nucl and Cyto fractions were measured by qPCR. The ratio of Nucl to Cyto mRNA was calculated and plotted. Points indicate individual biological repeats and error bars show the mean ± SEM. Significance was determined by multiple paired t test.

## Discussion

Coronaviruses encode several accessory proteins which are dispensable for *in vitro* replication but are presumably important for replication *in vivo* and disease pathogenicity [7]. This could make them good therapeutic targets. These proteins are not well characterised for SARS-CoV-2, so we decided to investigate their function. To do this, we generated a panel of lentiviral VLPs encoding each individual accessory protein with which to introduce each protein into various cells for defined functional assays. Interestingly however, we were repeatedly unable to generate VLPs encoding ORF6 (Fig 1A). We therefore determined how ORF6 inhibits VLP production. We showed that ORF6 not only supresses expression of the HIV-1 structural protein Gag, which is required for VLP synthesis, but it also reduced MLV Gag (S1 Fig) and mApple expression (Fig 1D), implying that ORF6 blocks protein expression more broadly, in a sequence-independent manner.

Protein expression could have been inhibited via multiple mechanisms, however, ORF6 has been reported to interact with Rae1 and Nup98 which are involved in nuclear import and export suggesting that ORF6 may modulate mRNA nuclear export resulting in decreased protein expression. We confirmed that ORF6 from both SARS-CoV-2 and SARS-CoV-1 interacted with Rae1, and that this interaction was important for the block to protein expression as mutants that were unable to interact also failed to inhibit protein expression (Figs 2 and 1F). Furthermore, we showed directly that ORF6 did indeed inhibit mRNA export (Figs 3 and 4). Interestingly, the herpesvirus ORF10 protein and the VSV M protein also interact with the same region of Rae1 [13, 15, 22] and block mRNA nuclear export. VSV M broadly inhibits the loading of cellular mRNA to the Rae1-Nup98 complex, while herpesvirus ORF10 can still form a complex with cellular mRNAs. Here, ORF6 only interacted with Nup98 indirectly through Rae1, similar to herpesvirus ORF10. ORF6 has been reported to dislocate Rae1 and Nup98 from the NPC, but whether this mislocalisation causes the mRNA export block or is a consequence of mRNA export disruption is not known [19].

Blocking cellular mRNA nuclear export, thereby reducing translation of cellular proteins, would be advantageous for viral replication as SARS-CoV-2 replicates in the cytoplasm and would therefore be able to exploit the limited cellular translational machinery in favour of viral translation (Fig 6A). In addition, by blocking mRNA export, ORF6 can also affect other cellular processes beyond export, as inhibiting the translation of key cellular proteins will modulate their downstream processes. Excitingly, using RNAseq, we have revealed that ORF6 can prevent the nuclear export of mRNAs encoding antiviral factors (Fig 4), both after IFN treatment, for example IFITM2 and ZNFX1, and more provocatively, in untreated cells. This implies that ORF6 can modulate the basal, steady state level of immunity in cells. Indeed, DDX58(RIG-I) and ZNFX1, which were enriched in the nucleus following both ORF6 expression in untreated cells (Fig 4) and during SARS-CoV-2 infection (Fig 5), are key pattern recognition receptors (PRRs) that detect viral dsRNA [23, 24]. ZNFX1 has been postulated to be involved in the very early stages of IFN signalling, as one of the earliest viral sensors, due to its higher constitutive expression compared to both MDA5 and RIG-I [23]. Downregulation of these sensors would prevent the induction of an antiviral state and allow SARS-CoV-2 to replicate undetected (Fig 6B). The constitutively expressed transcription factor, IRF1, which helps regulate basal expression of antiviral factors like BST2 and RIG-I [25, 26] was also targeted by ORF6. IRF1 mRNA is dependent on Nup98 and Rae1 for export and its corresponding protein has a relatively short half-life of around 20-40 mins [16, 27, 28]. Inhibiting IRF1 mRNA export would therefore have major consequences for the induction of innate immunity [16, 29, 30]. Interestingly, ORF6 from SARS-CoV-1 is thought to be packaged into virions and, thus, the protein is present in the cell as soon as virions infect. Incoming ORF6 could therefore downregulate PRRs before viral replication generates dsRNA intermediates, impeding the earliest antiviral warning system whilst also restricting the expression of further antiviral factors that should induce an antiviral state in the cell [31, 32]. It would be interesting to know if SARS-CoV-2 ORF6 is also packaged into virions.

**Fig 6.**
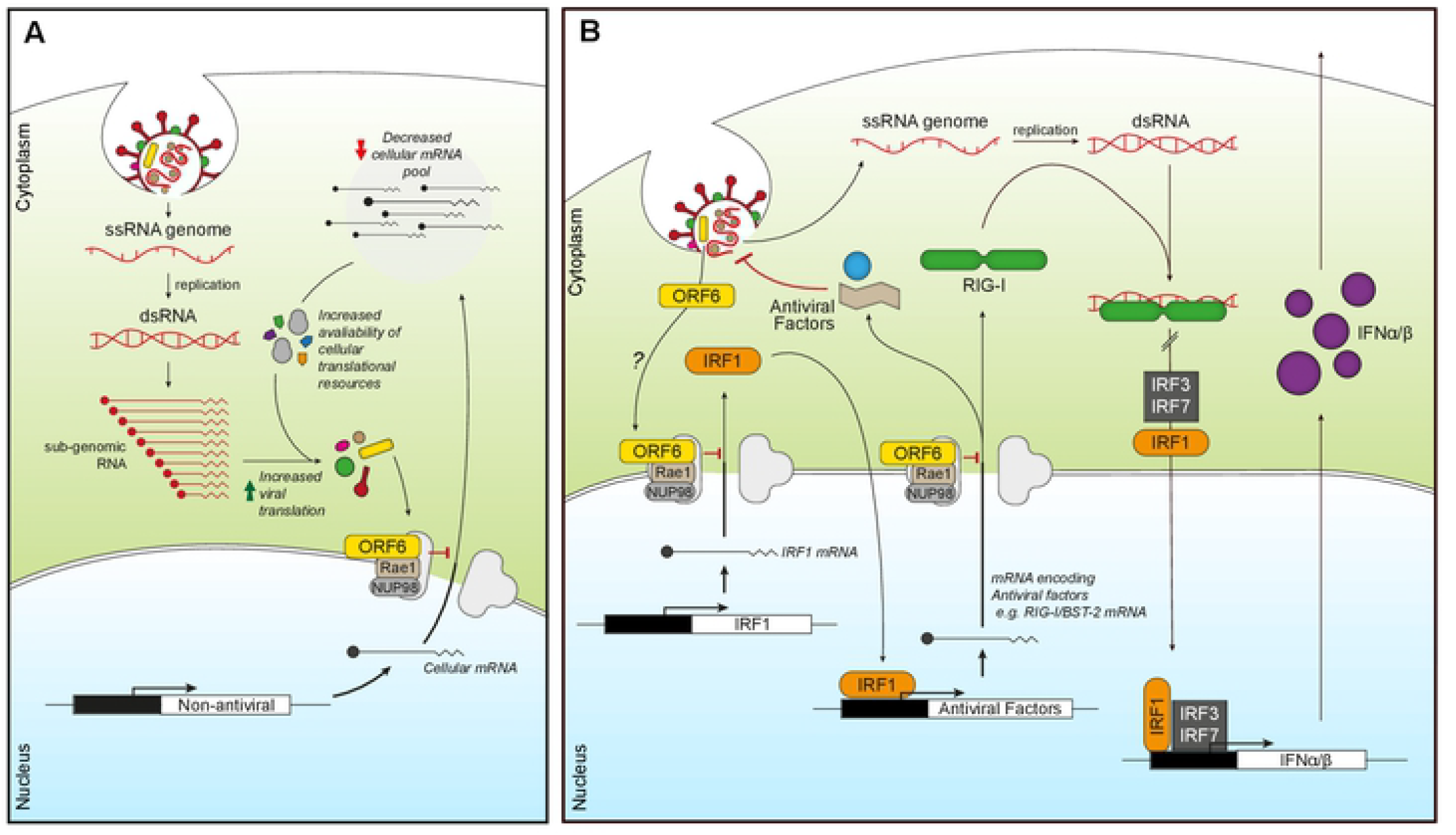
ORF6 inhibits mRNA export to favour viral translation and supress innate signalling. (A) Incoming or expressed ORF6 (yellow rectangles) binds to the Nup98-Rae1 complex (grey/brown) blocking the export of cellular mRNA (black lines). This subsequently reduces the cellular mRNA pool, increasing the availability of cellular translational machinery for viral translation as well as decreasing the expression of cellular proteins. (B) Incoming or expressed ORF6 (yellow) binds to the Nup98-Rae1 complex (grey/brown), inhibiting export of cellular mRNA encoding IRF1 (orange). This prevents the translation of IRF1, blocking IRF1 regulation of the transcription of additional steady state antiviral factors, like RIG-I and BST-2. ORF6 also inhibits nuclear export of mRNA encoding RIG-I (green), preventing detection of viral dsRNA produced during coronavirus replication. This helps reduce IRF1/3/7 activity and subsequent transcription of IFNα/β (purple), which prevents IFNα/β inducing an antiviral state in an autocrine and paracrine manner.

ORF6 from SARS-CoV-2 and SARS-CoV-1 have been shown to inhibit ISG and type-I IFN expression by preventing STAT1/2 and IRF3 nuclear translocation [9, 10, 14, 33, 34]. Another recent study used immunofluorescence to show that mRNA accumulated in the nucleus of ORF6 expressing cells [18]. Here, we show that ORF6 alone, or in the context of a viral infection, can inhibit nuclear export of a broad range of mRNA, including mRNA of antiviral proteins and transcription factors, which could have a downstream effect on ISG expression. Indeed, IRF1 regulates the expression of a range of ISGs by acting as a positive regulator of the JAK/STAT signalling pathway (Fig 6) [35]. Although ORF6 clearly inhibits mRNA nuclear export, the multi-functional SARS-CoV-2 protein, NSP1, has also recently been reported to inhibit mRNA export by blocking NXF1 docking to the NPC [17]. This redundancy in targeting mRNA export suggests that blocking this pathway gives the virus a clear survival advantage [36]. Further work will investigate how ORF6 and NSP1 complement each other and whether this can be exploited in new therapeutic strategies directed against Sarbecoviruses [17, 37].

## Materials and Methods

### Cell lines

HEK293T, HeLa, Mus dunni tail fibroblast (MDTF) and Vero cell lines were maintained in Dulbecco’s modified Eagle medium (DMEM, Thermo Fisher), supplemented with 10% heat-inactivated foetal bovine serum (FBS, Biosera) and 1% Penicillin/Streptomycin (Sigma). Cells were grown in a humified incubator at 37°C and 5% CO_2_. Stock cells were tested for mycoplasma contamination and authenticated by short-tandem repeat (STR) profiling.

### Plasmids and site-directed mutagenesis

SARS-CoV-2 ORF accessory proteins were expressed in cells via transfection or transduction with retroviral VLPs. The plasmids used to generate HIV-1 and MLV VLPs for transduction of ORFs, pCMVΔ8.91 and pHIT60 respectively, and pVSV-G have been described before [38, 39]. For expression of each SARS-CoV-2 ORF accessory protein, two different plasmids were used. For the initial experiments on Gag expression (Figs 1A-C and S1A and B Figs) and ORF6 co-expression with Rae1 (Fig 2A), a pLVX-IRES-Puro vector encoding an individual ORF was used. The ORF genes were initially synthesised by GeneArt (Thermo Fisher) and first cloned into a pTriEX-6 plasmids using XmaI and SacI restriction sites. The pLVX-StrepII-ORF-IRES-Puro plasmids were then cloned using the NEBuilder HiFi DNA Assembly Cloning Kit (NEB) from the pTriEX-6 plasmids, using the primers listed in Table 1. For the remaining assays that required ORF detection or immunoprecipitation, a pLVX-EF1α-SARS-CoV-2-ORF-2xStrep/2xStrep-ORF -IRES-Puro encoding one of the nine ORFs or pLVX-EF1α-eGFP-2xStrep-IRES-Puro plasmid was used, referred to as twin-strep-tag. These plasmids were kindly gifted from Nevan Krogan (Addgene plasmid #141383-4, #141387-90, #141392-95); http://n2t.net/addgene:141383-4, 141387-90, 141392-95; RRID:Addgene_141383-4, 141387-90, 141392-95) [5]. To generate the pLVX-EF1α-CoV-1-ORF6-2xStrep-IRES-Puro plasmid, the ORF6(CoV-1)-2XStrep gene was synthesised by GeneArt and cloned in using the EcoRI and BamHI restriction sites. ORF6 mutations were introduced into the pLVX-StrepII-SARS-CoV-2-ORF6-IRES-Puro and pLVX-EF1α-SARS-CoV-2-ORF6-2xStrep-IRES-Puro plasmids using the QuickChange II-XL site-directed mutagenesis kit (Agilent) according to the manufacturer’s instructions using SDM primers listed below. The following plasmids were sourced from Addgene, psfGFP-N1 (Addgene, #54737) and pmApple-N1 (Addgene, #54567). pCMVsport6Rae1 (MGC:117333) was purchased from Horizon Discovery Biosciences Limited. Nup98 was amplified from a HeLa cDNA library from Clontech/Takara. Affinity tags were introduced by NEBuilder HiFi DNA Assembly Cloning Kit (NEB) into the expression vectors encoding Rae1 and Nup98 using primers in Table 1. Sequences were verified by Sanger sequencing (Source Bioscience).

**Table 1.**
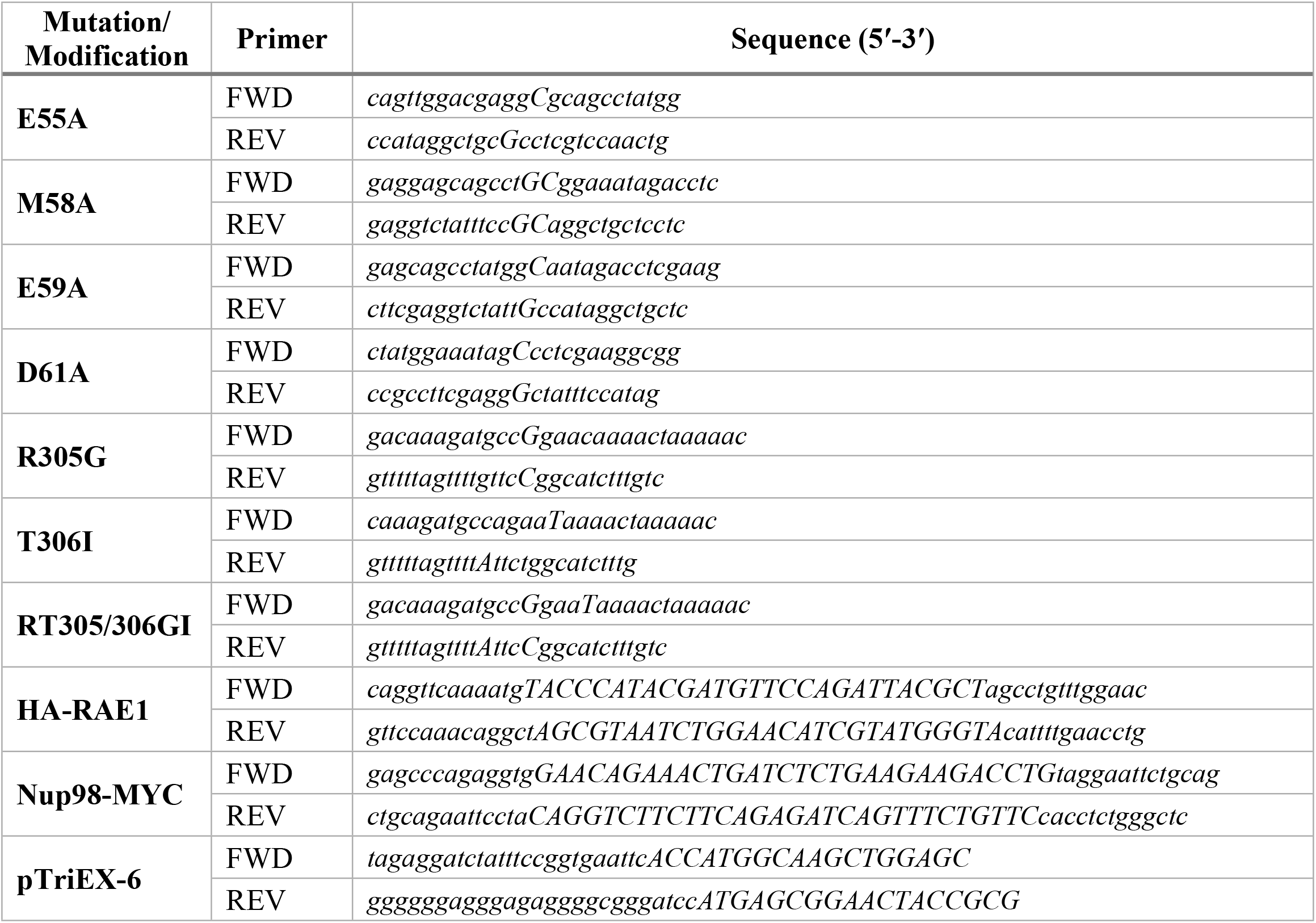
Sequences of Primers used for cloning.

### Virus-like particles (VLP) production

HIV-1 and MLV virus-like particles were generated by co-transfecting HEK293T cells with plasmids, pCMVΔ8.91 (HIV-1) or pHIT60 (MLV) and pVSV-G using lipofectamine 2000 (11668019, Invitrogen). For generating GFP reporter VLPs, pCSGW (HIV-1) or pczCFG2fEGFPf (MLV) were also added [39, 40]. The plasmids were used at a ratio of 1:1:1. After 16 hrs, cells were treated with 10 mM sodium butyrate for 8 hrs, and supernatant containing VLPs harvested 24 hrs later. To measure viral titres, HeLa cells were transduced with serially diluted supernatant containing the VLPs. After 72 hrs, cells were selected with media containing puromycin (1 μg/ml) for 10 days, after which cells were washed, fixed, and stained with methanol containing crystal violet. Colonies were counted and viral titre calculated.

### Immunoblotting

Cells were lysed in ice cold radioimmunoprecipitation assay (RIPA) buffer (Thermo Fisher) or IP buffer (50 mM Tris-HCl, pH 7.4 at 4 °C, 150 mM NaCl, 1 mM EDTA supplemented with 0.5% Nonidet P40) containing protease inhibitor cocktail (Roche) and DNAse (#88701, Thermo Fisher). Protein lysates were boiled at 95 °C for 5 mins in 6x SDS sample buffer (Thermo Scientific). Proteins were separated on 4-12% Bis-Tris SDS-PAGE gels (Thermo Fisher) or 4-20% mini-PROTEAN TGX (Bio-Rad) and transferred to a PVDF membrane. Primary antibodies used were: anti-HIV-1 p24 (ARP432, CFAR), anti-GFP (sc-9996, SCBT), anti-HA (C29F4, CST), anti-Strep-tag (34850, Qiagen), anti-HSP90 (4874, CST), anti-Histone H3 (14269, CST), anti-alpha-Tubulin (VMA00051, BIO-RAD), anti-Lamin B1 (66095-1-Ig, Proteintech), anti-Nup98 (C39A3, CST), anti-Rae1 (ab124783, Abcam), anti-pSTAT(Y701) (D4A7, CST), anti-ORF6/9b (MRC PPU, Dundee University). Secondary antibodies include, anti-mouse (61-6520, Thermo Fisher), anti-sheep (ab6747, Abcam) and anti-rabbit (31460, Thermo Fisher) HRP-conjugated; anti-mouse IRDye 680RD/800CW (926-68072/926-32212, Li-Cor) and anti-rabbit IRDye 700CW/800CW (926-68073/926-32213, Li-Cor). Blots were imaged on a Chemidoc MP imaging system (Bio-Rad) or Odyssey Clx imaging system (LICOR).

### Immunoprecipitation

HEK293T cells were grown in 10 cm^2^ culture plates transfected and incubated for 16 hrs after which media was replaced. After a further 8 hrs, cells were harvested in Pierce IP lysis buffer (Thermo Fisher), supplemented with protease inhibitor cocktail and Pierce DNAse (Thermo Fisher). Twin-strep fusion proteins were immunoprecipitated with MagStrep “type3” XT beads (IBA) as per the manufacturer’s instructions. Proteins were eluted with biotin (50 mM) and boiled in 6x SDS sample buffer.

### Subcellular fractionation

HEK293T cells were transfected with psfGFP-N1 and either treated with LMB (Merck) at 5nM; or co-transfected with plasmids expressing pLVX-EF1α-ORF6, ORF6(M58A) or ORF9b [5]. At the indicated times post-transfection, 1×10^6^ cells were harvested and processed with the PARIS kit (Life Technologies) following the manufacturer’s instructions. Parallel, samples were lysed as total cell lysates or were fractionated into cytoplasm and nuclear fractions. Each sample was then split in two: half was used for protein analysis and half was used for RNA extraction. Protein content in the total cell lysate and in the fractions was measured by BCA assay (Thermo Fisher) using a FLUOstar Omega plate reader (BMG Labtech). An equivalent amount of each fraction compared to the total cell lysate was analysed by immunoblotting.

### Quantitative PCR

To remove possible DNA contamination, extracted RNA from the nuclear and cytoplasmic fractions was treated with DNase using the DNA-free kit (Invitrogen) and following the manufacturer’s instructions. Then, 100ng of RNA was converted to cDNA for 1h at 37 °C using the Omniscript RT kit (Qiagen) and random primers (Promega). Quantitative PCR (qPCR) analysis was performed in TaqMan Fast Advanced master mix (Thermo Fisher) with 900nM primers and 250nM probes, or 1X Taqman gene expression assays. The reactions were performed on a 7500 fast real-time PCR system (Applied Biosystems) using standard cycling conditions: 50 °C for 2min, 95 °C for 10min followed by 40 cycles of 95 °C for 15s and 60 °C for 1min. To calculate DNA copy numbers, standard curves were generated from serial dilutions of the indicated cDNA in water. The following primers and probes were used; sfGFP: *for* 5’-GCGCACCATCAGCTTCAAGG, *rev* 5’-GTGTCGCCCTCGAACTTCAC and *probe* 5’-*FAM-*CGGCACCTACAAGACCCGCGC*-TAMRA*.

### mRNAseq

The quality of extracted RNA was checked by Agilent TapeStation 42000 on Agilent RNA ScreenTape (Agilent technologies) before proceeding to library preparation. Only samples with a RINe of 7 or higher were used for Library preparation. RNA-seq libraries were generated using the NEBNext Ultra Directional RNA Library Prep kit for Illumina with NEBNext® Poly(A) mRNA Magnetic Isolation Module (both New England BioLabs). Samples were adjusted to 400 ng total RNA and spiked with diluted 1/100 ERCC Spike-In Mix (4456740, Invitrogen) and Drosophila melanogaster, embryo Poly A+ RNA, 5 μG (636224-Takara) using 1 μl from each. PCR Enrichment of Adaptor Ligated DNA was run with 12 cycles and SPRI beads (Beckman) were used for library clean up. Agilent DNA Screen tapes (Agilent Technologies) were used to check library size and quality. Samples were sequenced on an Illumina Novaseq 6000 platform with 180 pM loading concentration in single-end run using 100-bp reads. Enriched GO terms (Biological process) were filtered and mapped using the REViGO tool available at http://revigo.irb.hr/ [41]. The raw RNAseq data from this study can be accessed from ENA via accession PRJEB49943.

### mRNAseq Data Analysis

A custom genome joining the Human (Ensembl GRCh38.100) and D. melanogaster (Ensembl BDGP6.104) genomes was prepared, and reads were trimmed, aligned, and counts produced using the nf-core rnaseq pipeline [42]. Normalisation and differential expression analyses were conducted within DESeq2 [43]: normalisation factors for gene-level read counts were produced using counts for the D. melanogaster spike-in controls and differential gene expression calculations were subsequently performed for Human genes alone using the pre-calculated size factors. Independent hypothesis weighting was conducted to optimise the power of p value filtering using the IHW package [44] and Log2 fold-change shrinkage was performed using ashr package [45]. Result tables were subsequently filtered on (IHW-adjusted) q<0.05.

### SARS-CoV-2 Infections

The following viruses were used in the infections: The Wuhan-like reference isolate (referred to as the Eng/2) was the hCoV-19/England/02/2020, obtained from the Respiratory Virus Unit, Public Health England, UK (GISAID EpiCov accession EPI_ISL_407073). The B.1.1.7 (Alpha) isolate was the hCoV-19/England/204690005/2020, which carries the D614G, Δ69– 70, Δ144, N501Y, A570D, P681H, T716I, S982A, and D1118H mutations [46], obtained from Public Health England (PHE), UK, through Prof. Wendy Barclay, Imperial College London, London, UK. The B.1.351 (Beta) virus isolate was the 501Y.V2.HV001, which carries the D614G,L18F,D80A, D215G, Δ242-244, K417N, E484K, N501Y, A701V mutations, and was kindly provided by Prof. Alex Sigal and Prof. Tulio de Oliveira [47], sequencing of viral isolates received identified the Q677H and R682W mutations at the furin cleavage site in approximately 50% of the genomes, which was maintained upon passage in cell culture. The B.1.617.2 (Delta) isolate was MS066352H (GISAID accession number EPI_ISL_1731019), which carries the T19R, K77R, G142D, Δ156-157/R158G, A222V, L452R, T478K, D614G, P681R, D950N, and was kindly provided by Prof. Wendy Barclay, Imperial College London, London, UK through the Genotype-to-Phenotype National Virology Consortium (G2P-UK). Viruses were titrated by plaque assay on Vero E6 cells. Serial ten-fold dilutions of virus are incubated for 40 mins at RT on a well of confluent Vero E6 cells. The virus inoculum is then removed, and the cells overlayed with a semisolid overlay containing 1.2% Avicel in MEM. Cells are incubated for two days at 37 °C, 5 % CO_2_ after which the overlay is removed. Cells were fixed and stained with 4 % paraformaldehyde containing 0.2 % toluidine blue for at least 30 mins. For the infections, Vero E6 cells were seeded 24 hrs before use to achieve 80 - 90% confluency. Cells were washed, medium was removed from each well and 1 ml of serum-free DMEM added. The 1 ml of serum-free DMEM was removed and 500 µl virus diluted in virus growth medium was added to achieve an MOI of ≥1. Plates were incubated for approx. 30 mins at RT and then 1 ml virus growth medium was added to each well. Plates were incubated for 24 hrs before RNA isolation and qPCR as described above using Taqman gene expression assays (Thermo Fisher) as follows: FAM222A (Hs00757936_m1), SPOCK1 (Hs00270274_m1), SERTAD1 (Hs00203547_m1), ZNFX1 (Hs01105231_m1), TRIM38 (Hs00197164_m1), RIG-I (Hs01061433_m1), GAPDH (Hs02786624_g1) and IRF1 (Hs00971964_g1).

### Immunofluorescence

HEK293T cells were grown on glass coverslips and transfected or infected with SARS-CoV-2. Cells were fixed in 4% PFA in PBS for 10 mins at RT. Cells were then permeabilised with 0.1% triton in PBS and blocked in 3% BSA in PBS. Cells were labelled with the following primary antibodies, anti-Nup98 or anti-ORF6 diluted in 4% BSA in PBS for 1 hour at RT. Cells were then incubated with the following secondary antibodies, donkey anti-rabbit-AF568 (ab175692, Abcam) and donkey anti-sheep AF568 (A-21099, Invitrogen) diluted in 4% BSA in PBS for 1 hour at RT. Coverslips were washed and mounted on glass coverslips with ProLong Gold Antifade Mountant with DAPI (Thermo Fisher) and imaged with the Leica SP5 microscope and 100X 1.3NA oil immersion objective (Leica).

### Statistics

Statistical analysis was performed using GraphPad Prism 9 software. (*, P < 0.05; **, P < 0.01; ***, P < 0.001; ****, P < 0.0001, ns – not significant).

## Acknowledgements

We thank Nevan Krogan for gifting the twin-strep-tagged ORF protein plasmids via Addgene (Addgene plasmid #141383-4, #141387-90, #141392-95) and Wendy Barclay, Prof. Alex Sigal and Prof. Tulio de Oliveira for providing SARS-CoV-2 viral isolates. We are extremely grateful to Jimena Perez-Lloret, Robert Goldstone and Deb Jackson from the Crick ASF STP for running the RNAseq experiment. For the purpose of Open Access, the authors have applied a CC BY public copyright licence to any Author Accepted Manuscript version arising from this submission.

## Funding

This work was supported by the Francis Crick Institute, which receives its core funding from Cancer Research UK (FC001042, to KB and FC001162, to JS), the UK Medical Research Council (FC001042, to KB and FC001162, to JS) and the Wellcome trust (FC001042, to KB and FC001162, to JS) and by the MRC Genotype-to-Phenotype (G2P) UK National Virology Consortium grant to KB (MR/W005611/1) and by a Wellcome Trust Investigator Award to JS (108012/Z/15/Z). The funders had no role in study design, data collection and analysis, decision to publish, or preparation of the manuscript.

## Supporting information

**S1 Fig. SARS-CoV-2 ORF6 Inhibits MLV Gag expression**.

(A, B) To generate MLV VLPs, HEK293T cells were transfected with plasmids encoding Gag-Pol(MLV), VSV-G and either increasing amounts of pLVX-StrepII-ORF6 or pLVX-StrepII-ORF9b as well as a GFP reporter plasmid, pczCFG2fEGFPf. As a control, VLPs were generated with an empty pLVX vector. (A) Transfected cells were lysed and analysed by SDS-PAGE and immunoblotting for Gag(MLV), ORF6 or ORF9b and HSP90. (B) MDTF cells were infected with increasing amounts of MLV VLPs. Three days post infection, the percentage of GFP positive cells was determined by flow cytometry. Graph shows the mean and range of two biological repeats. (C) HEK293T cells were transfected with the pLVX-EF1α plasmid encoding either ORF6(CoV-1), ORF6(CoV-2) or the indicated ORF6 mutant as well as a plasmid encoding mApple. After 48h, cells were fixed and the mApple MFI measured by flow cytometry. The fold change (FC) in MFI is plotted relative to the control for independent biological repeats. Error bars show SEM. Transfected cell lysates were analysed by SDS-PAGE and immunoblotting with anti-ORF6.

**S2 Fig. ORF6 interacts with endogenous Rae1**.

(A) HEK293T cells were transfected with plasmids encoding Twin-Strep-tagged GFP, ORF6(CoV-2) or ORF6(CoV-1) and HA-Rae1. After 24h, tagged proteins were immunoprecipitated with MagStrep beads and proteins eluted with biotin. Input lysates and eluate were separated by SDS-PAGE and analysed by immunoblotting for Strep, Nup98 and HA. (B) HEK293T cells were transfected with the pLVX-EF1α plasmid encoding Twin-Strep-tagged GFP or ORF6. After 24h, tagged proteins were immunoprecipitated with MagStrep beads and proteins eluted with biotin. Input lysates and eluate were separated by SDS-PAGE and analysed by immunoblotting for endogenous Rae1 and anti-Strep.

**S3 Fig. ORF6 inhibits GFP mRNA export in a Rae1-dependent manner**.

(A-D) HEK293T cells were transfected with a plasmid expressing GFP or left untransfected (mock) and either treated with 5 nM Leptomycin B (LMB) or left untreated. After 16h, cells were either harvested as a total cell lysate (T) or fractionated into Nucl (N) and Cyto (C) fractions. Total cell lysates and cellular fractions were divided into two for either protein analysis (B) or RNA extraction (C,D). (A) Low magnification fluorescent images showing GFP expression in transfected cells before harvest. (B) Protein levels were quantified by BCA assay, proportional amounts of the fractions related to the total cell lysate were analysed by immunoblotting for GFP, HSP90 (cytoplasmic marker) and Histone H3 (nuclear marker). (C) RNA from Nucl and Cyto fractions was converted into cDNA and GFP mRNA copy numbers measured by qPCR. (D) The ratio of Nucl to Cyto GFP mRNA was calculated and plotted for individual biological repeats. Error bars show the mean ± SEM. (E-H) HEK293T cells were co-transfected with plasmids encoding either ORF6, ORF6(M58A) or ORF9b and GFP, or left untransfected (mock). After 32h, cells were either harvested as a total cell lysate (T) or fractionated into Nucl (N) and Cyto (C) fractions. Total cell lysates and cellular fractions were divided into two for either protein analysis (F) or RNA extraction (G,H). (E) Low magnification fluorescent images showing GFP expression in transfected cells before harvest. (F) Protein levels were quantified by BCA assay, proportional amounts of the fractions related to the total cell lysate were analysed by immunoblotting for Strep, GFP, HSP90 (cytoplasmic marker) and Histone H3 (nuclear marker). (G) RNA from Nucl and Cyto fractions was converted into cDNA and GFP mRNA copy numbers measured by qPCR. (H) The ratio of Nucl to Cyto GFP mRNA was calculated and plotted for individual biological repeats. Error bars show the mean ± SEM. Significance was calculated by one-way ANOVA with Turkey’s post hoc test, *p<0.05, **p<0.01, ns – non-significant.

**S4 Fig. ORF6 inhibits the nuclear export of cellular mRNA**.

(A) The cell fractionations used in Fig 3A, and B were analysed by immunoblotting with anti-Strep to label ORF6 and GFP, anti-pSTAT(Y701) to confirm activation of the type-I IFN signalling cascade and anti-HSP90 (cytoplasmic marker) or anti-Lamin B1 (nuclear marker) to confirm successful fractionation. The analysis of three biological repeats is shown. (B) From the mRNAseq data in Fig 3. The log2-fold change (Log2-FC) in mRNA abundance was compared between the Nucl and Cyto fractions for both GFP and ORF6 expressing cells, without (upper panels) and with IFN treatment (lower panels). The log-2FC was weighted against the adjusted *p*-value (shrunken Log2-FC) to show mRNAs that are significantly enriched (GFP; black, ORF6; yellow) in either the cytoplasm or nucleus. The number of mRNAs significantly enriched are shown. The mRNA count was normalised and averaged between three biological repeats.

**S5 Fig. ORF6 inhibits the nuclear export of IFN-upregulated mRNA**.

(A) From the mRNAseq data in Fig 3. The Log2-FC in mRNA abundance was compared between cells treated with IFN or left untreated in both the Cyto (left panel) and Nucl (right panel) fractions. The log-2FC was weighted against the adjusted *p*-value (shrunken Log2-FC) to show mRNA species that are significantly upregulated or downregulated (blue). The number of mRNAs upregulated is shown. These were designated as Interferon upregulated genes (IUGs). (B) Venn diagram showing the number of IUGs significantly upregulated in Cyto. and Nucl. fractions. (C) GO ontology enrichment analysis of significantly upregulated IUGs from A. in both the Cyto and Nucl fractions. (D) Venn diagram showing the number of IUGs significantly downregulated (red) or upregulated (green) by ORF6 in Cyto. and Nucl. fractions, as described in Fig 4A. (E) From the mRNAseq data in Fig 3. The log2-fold change (Log2-FC) in mRNA abundance was compared between ORF6 and GFP expressing cells in the Cyto. and Nucl fractions, without (upper panel) and with IFN treatment (lower panel). This log2-FC was weighted against the adjusted *p*-value (shrunken Log2-FC) to identify significantly upregulated/downregulated mRNAs (in yellow), with the number of mRNAs shown. The mRNA count was normalised and averaged between the three biological repeats.

**S6 Fig. SARS-CoV-2 inhibits the export of cellular mRNA**

(A) Vero cells that were inoculated with SARS-CoV-2(beta) in Fig 5A were fixed, permeabilised and analysed by immunofluorescence with anti-Nup98, anti-ORF6 and DAPI. (B-D) Cell lysates from Fig 5B were analysed by immunoblotting for Strep to label ORF6, Nsp2, HSP90 (Cytoplasmic marker) and Histone H3 (Nuclear marker) to confirm successful fractionation. The analysis of three biological repeats are shown. (E) The plot shown in Fig 3B (right panel) highlighting the enrichment of the mRNA encoding the four non-antiviral proteins that were quantified by qPCR in Fig 5B.

## Notes

### Competing Interest Statement

The authors have declared no competing interest.

## References

1. Lu R, Zhao X, Li J, Niu P, Yang B, Wu H, et al. Genomic characterisation and epidemiology of 2019 novel coronavirus: implications for virus origins and receptor binding. The Lancet. 2020;395(10224):565–74. doi: 10.1016/s0140-6736(20)30251-8.

2. Xu Z, Shi L, Wang Y, Zhang J, Huang L, Zhang C, et al. Pathological findings of COVID-19 associated with acute respiratory distress syndrome. The Lancet Respiratory Medicine. 2020;8(4):420–2. doi: 10.1016/s2213-2600(20)30076-x.

3. The species Severe acute respiratory syndrome-related coronavirus: classifying 2019-nCoV and naming it SARS-CoV-2. Nature Microbiology. 2020;5(4):536–44. doi: 10.1038/s41564-020-0695-z.

4. Zhou P, Yang X-L, Wang X-G, Hu B, Zhang L, Zhang W, et al. A pneumonia outbreak associated with a new coronavirus of probable bat origin. Nature. 2020;579(7798):270–3. doi: 10.1038/s41586-020-2012-7.

5. Gordon DE, Jang GM, Bouhaddou M, Xu J, Obernier K, White KM, et al. A SARS-CoV-2 protein interaction map reveals targets for drug repurposing. Nature. 2020;583(7816):459–68. doi: 10.1038/s41586-020-2286-9.

6. Schaecher SR, Pekosz A. SARS Coronavirus Accessory Gene Expression and Function. Molecular Biology of the SARS-Coronavirus: Springer Berlin Heidelberg; 2010. p. 153–66.

7. Liu DX, Fung TS, Chong KK-L, Shukla A, Hilgenfeld R. Accessory proteins of SARS-CoV and other coronaviruses. Antiviral Research. 2014;109:97–109. doi: 10.1016/j.antiviral.2014.06.013.

8. Kopecky-Bromberg SA, MartíNez-Sobrido L, Frieman M, Baric RA, Palese P. Severe Acute Respiratory Syndrome Coronavirus Open Reading Frame (ORF) 3b, ORF 6, and Nucleocapsid Proteins Function as Interferon Antagonists. Journal of Virology. 2007;81(2):548–57. doi: 10.1128/jvi.01782-06.

9. Frieman M, Yount B, Heise M, Kopecky-Bromberg SA, Palese P, Baric RS. Severe Acute Respiratory Syndrome Coronavirus ORF6 Antagonizes STAT1 Function by Sequestering Nuclear Import Factors on the Rough Endoplasmic Reticulum/Golgi Membrane. Journal of Virology. 2007;81(18):9812–24. doi: 10.1128/jvi.01012-07.

10. Miorin L, Kehrer T, Sanchez-Aparicio MT, Zhang K, Cohen P, Patel RS, et al. SARS-CoV-2 Orf6 hijacks Nup98 to block STAT nuclear import and antagonize interferon signaling. Proceedings of the National Academy of Sciences. 2020;117(45):28344–54. doi: 10.1073/pnas.2016650117.

11. Bharathi A, Ghosh A A. Whalen W, Ho Yoon J, Pu R, Dasso M, et al. The human RAE1 gene is a functional homologue of Schizosaccharomyces pombe rae1 gene involved in nuclear export of Poly(A)+ RNA. Gene. 1997;198(1-2):251–8. doi: 10.1016/s0378-1119(97)00322-3.

12. Pritchard CEJ, Fornerod M, Kasper LH, Van Deursen JMA. RAE1 Is a Shuttling mRNA Export Factor That Binds to a GLEBS-like NUP98 Motif at the Nuclear Pore Complex through Multiple Domains. Journal of Cell Biology. 1999;145(2):237–54. doi: 10.1083/jcb.145.2.237.

13. Faria PA, Chakraborty P, Levay A, Barber GN, Ezelle HJ, Enninga J, et al. VSV Disrupts the Rae1/mrnp41 mRNA Nuclear Export Pathway. Molecular Cell. 2005;17(1):93–102. doi: 10.1016/j.molcel.2004.11.023.

14. Her L. Inhibition of Ran Guanosine Triphosphatase-Dependent Nuclear Transport by the Matrix Protein of Vesicular Stomatitis Virus. Science. 1997;276(5320):1845–8. doi: 10.1126/science.276.5320.1845.

15. Von Kobbe C, Van Deursen Jma, Rodrigues JP, Sitterlin D, Bachi A, Wu X, et al. Vesicular Stomatitis Virus Matrix Protein Inhibits Host Cell Gene Expression by Targeting the Nucleoporin Nup98. Molecular Cell. 2000;6(5):1243–52. doi: 10.1016/s1097-2765(00)00120-9.

16. Satterly N, Tsai P-L, Van Deursen J, Nussenzveig DR, Wang Y, Faria PA, et al. Influenza virus targets the mRNA export machinery and the nuclear pore complex. Proceedings of the National Academy of Sciences. 2007;104(6):1853–8. doi: 10.1073/pnas.0610977104.

17. Zhang K, Miorin L, Makio T, Dehghan I, Gao S, Xie Y, et al. Nsp1 protein of SARS-CoV-2 disrupts the mRNA export machinery to inhibit host gene expression. Science Advances. 2021;7(6):eabe7386. doi: doi:10.1126/sciadv.abe7386.

18. Addetia A, Lieberman NAP, Phung Q, Hsiang T-Y, Xie H, Roychoudhury P, et al. SARS-CoV-2 ORF6 Disrupts Bidirectional Nucleocytoplasmic Transport through Interactions with Rae1 and Nup98. mBio. 2021;12(2):e00065–21. doi: 10.1128/mBio.00065-21. PubMed PMID: 33849972.

19. Kato K, Ikliptikawati DK, Kobayashi A, Kondo H, Lim K, Hazawa M, et al. Overexpression of SARS-CoV-2 protein ORF6 dislocates RAE1 and NUP98 from the nuclear pore complex. Biochemical and Biophysical Research Communications. 2021;536:59–66. doi: 10.1016/j.bbrc.2020.11.115.

20. Ren Y, Seo H-S, Blobel G, Hoelz A. Structural and functional analysis of the interaction between the nucleoporin Nup98 and the mRNA export factor Rae1. Proceedings of the National Academy of Sciences. 2010;107(23):10406–11. doi: 10.1073/pnas.1005389107.

21. Watanabe M, Fukuda M, Yoshida M, Yanagida M, Nishida E. Involvement of CRM1, a nuclear export receptor, in mRNA export in mammalian cells and fission yeast. Genes to Cells. 1999;4(5):291–7. doi: 10.1046/j.1365-2443.1999.00259.x.

22. Gong D, Kim YH, Xiao Y, Du Y, Xie Y, Lee KK, et al. A Herpesvirus Protein Selectively Inhibits Cellular mRNA Nuclear Export. Cell Host & Microbe. 2016;20(5):642–53. doi: 10.1016/j.chom.2016.10.004.

23. Wang Y, Yuan S, Jia X, Ge Y, Ling T, Nie M, et al. Mitochondria-localised ZNFX1 functions as a dsRNA sensor to initiate antiviral responses through MAVS. Nature Cell Biology. 2019;21(11):1346–56. doi: 10.1038/s41556-019-0416-0.

24. Yoneyama M, Kikuchi M, Natsukawa T, Shinobu N, Imaizumi T, Miyagishi M, et al. The RNA helicase RIG-I has an essential function in double-stranded RNA-induced innate antiviral responses. Nature Immunology. 2004;5(7):730–7. doi: 10.1038/ni1087.

25. Su Z-Z, Sarkar D, Emdad L, Barral PM, Fisher PB. Central role of interferon regulatory factor-1 (IRF-1) in controlling retinoic acid inducible gene-I (RIG-I) expression. Journal of Cellular Physiology. 2007;213(2):502–10. doi: 10.1002/jcp.21128.

26. Panda D, Gjinaj E, Bachu M, Squire E, Novatt H, Ozato K, et al. IRF1 Maintains Optimal Constitutive Expression of Antiviral Genes and Regulates the Early Antiviral Response. Front Immunol. 2019;10(1019). doi: 10.3389/fimmu.2019.01019.

27. Watanabe N, Sakakibara J, Hovanessian AG, Taniguchi T, Fujita T. Activation of IFN-βelement by IRF-1 requires a post-translational event in addition to IRF-1 synthesis. Nucleic Acids Research. 1991;19(16):4421–8. doi: 10.1093/nar/19.16.4421.

28. Nakagawa K, Yokosawa H. Degradation of transcription factor IRF-1 by the ubiquitin-proteasome pathway. European Journal of Biochemistry. 2000;267(6):1680–6. doi: 10.1046/j.1432-1327.2000.01163.x.

29. Michalska A, Blaszczyk K, Wesoly J, Bluyssen HAR. A Positive Feedback Amplifier Circuit That Regulates Interferon (IFN)-Stimulated Gene Expression and Controls Type I and Type II IFN Responses. Front Immunol. 2018;9:1135-. doi: 10.3389/fimmu.2018.01135. PubMed PMID: 29892288.

30. Pine R. Constitutive expression of an ISGF2/IRF1 transgene leads to interferon-independent activation of interferon-inducible genes and resistance to virus infection. Journal of Virology. 1992;66(7):4470–8. doi: 10.1128/jvi.66.7.4470-4478.1992.

31. Huang C, Peters CJ, Makino S. Severe Acute Respiratory Syndrome Coronavirus Accessory Protein 6 Is a Virion-Associated Protein and Is Released from 6 Protein-Expressing Cells. Journal of Virology. 2007;81(10):5423–6. doi: 10.1128/jvi.02307-06.

32. Sola I, Almazán F, Zúñiga S, Enjuanes L. Continuous and Discontinuous RNA Synthesis in Coronaviruses. Annual Review of Virology. 2015;2(1):265–88. doi: 10.1146/annurev-virology-100114-055218.

33. Li J-Y, Liao C-H, Wang Q, Tan Y-J, Luo R, Qiu Y, et al. The ORF6, ORF8 and nucleocapsid proteins of SARS-CoV-2 inhibit type I interferon signaling pathway. Virus Research. 2020;286:198074. doi: 10.1016/j.virusres.2020.198074.

34. Lei X, Dong X, Ma R, Wang W, Xiao X, Tian Z, et al. Activation and evasion of type I interferon responses by SARS-CoV-2. Nature Communications. 2020;11(1). doi: 10.1038/s41467-020-17665-9.

35. Pine R, Decker T, Kessler DS, Levy DE, Darnell JE, Jr. Purification and cloning of interferon-stimulated gene factor 2 (ISGF2): ISGF2 (IRF-1) can bind to the promoters of both beta interferon- and interferon-stimulated genes but is not a primary transcriptional activator of either. Mol Cell Biol. 1990;10(6):2448–57. Epub 1990/06/01. doi: 10.1128/mcb.10.6.2448-2457.1990. PubMed PMID: 2342456; PubMed Central PMCID: PMCPMC360601.

36. Finkel Y, Gluck A, Nachshon A, Winkler R, Fisher T, Rozman B, et al. SARS-CoV-2 uses a multipronged strategy to impede host protein synthesis. Nature. 2021;594(7862):240–5. doi: 10.1038/s41586-021-03610-3.

37. Stutz F. The interplay of nuclear mRNP assembly, mRNA surveillance and export. Trends in Cell Biology. 2003;13(6):319–27. doi: 10.1016/s0962-8924(03)00106-5.

38. Soneoka Y, Cannon PM, Ramsdale EE, Griffiths JC, Romano G, Kingsman SM, et al. A transient three-plasmid expression system for the production of high titer retroviral vectors. Nucleic Acids Research. 1995;23(4):628–33. doi: 10.1093/nar/23.4.628.

39. Bainbridge J, Stephens C, Parsley K, Demaison C, Halfyard A, Thrasher A, et al. In vivo gene transfer to the mouse eye using an HIV-based lentiviral vector; efficient long-term transduction of corneal endothelium and retinal pigment epithelium. Gene Therapy. 2001;8(21):1665–8. doi: 10.1038/sj.gt.3301574.

40. Zufferey R, Nagy D, Mandel RJ, Naldini L, Trono D. Multiply attenuated lentiviral vector achieves efficient gene delivery in vivo. Nature Biotechnology. 1997;15(9):871–5. doi: 10.1038/nbt0997-871.

41. Supek F, Bošnjak M, Škunca N, Šmuc T. REVIGO Summarizes and Visualizes Long Lists of Gene Ontology Terms. PLoS ONE. 2011;6(7):e21800. doi: 10.1371/journal.pone.0021800.

42. Ewels PA, Peltzer A, Fillinger S, Patel H, Alneberg J, Wilm A, et al. The nf-core framework for community-curated bioinformatics pipelines. Nature Biotechnology. 2020;38(3):276–8. doi: 10.1038/s41587-020-0439-x.

43. Love MI, Huber W, Anders S. Moderated estimation of fold change and dispersion for RNA-seq data with DESeq2. Genome Biology. 2014;15(12). doi: 10.1186/s13059-014-0550-8.

44. Ignatiadis N, Klaus B, Zaugg JB, Huber W. Data-driven hypothesis weighting increases detection power in genome-scale multiple testing. Nature Methods. 2016;13(7):577–80. doi: 10.1038/nmeth.3885.

45. Stephens M. False discovery rates: a new deal. Biostatistics. 2016:kxw041. doi: 10.1093/biostatistics/kxw041.

46. Brown JC, Goldhill DH, Zhou J, Peacock TP, Frise R, Goonawardane N, et al. Increased transmission of SARS-CoV-2 lineage B.1.1.7 (VOC 2020212/01) is not accounted for by a replicative advantage in primary airway cells or antibody escape. 2021.

47. Cele S, Gazy I, Jackson L, Hwa S-H, Tegally H, Lustig G, et al. Escape of SARS-CoV-2 501Y.V2 from neutralization by convalescent plasma. Nature. 2021;593(7857):142–6. doi: 10.1038/s41586-021-03471-w.

